# Dissecting Complexity: The Hidden Impact of Application Parameters on Bioinformatics Research

**DOI:** 10.1101/2022.12.20.521257

**Authors:** Mikaela Cashman, Myra B. Cohen, Alexis L. Marsh, Robert W. Cottingham

## Abstract

Biology is a quest; an ongoing inquiry about the nature of life. How do the different forms of life interact? What makes up an ecosystem? How does a tiny bacterium work? To answer these questions biologists turn increasingly to sophisticated computational tools. Many of these tools are highly configurable, allowing customization in support of a wide range of uses. For example, algorithms can be tuned for precision, efficiency, type of inquiry, or for specific categories of organisms or their component subsystems. Ideally, configurability provides useful flexibility. However, the complex landscape of configurability may be fraught with pitfalls. This paper examines that landscape in bioinformatics tools. We propose a methodology, SOMATA, to facilitate systematic exploration of the vast choice of application parameters, and apply it to three different tools on a range of scientific inquires. We further argue that the tools themselves are complex ecosystems. If biologists explore these, ask questions, and experiment just as they do with their biological counterparts, they will benefit by both finding improved solutions to their problems as well as increasing repeatability and transparency. We end with a call to the community for an increase in shared responsibility and communication between tool developers and the biologists that use them in the context of complex system decomposition.

## Introduction

Scientific investigation in biology has become increasingly dependent on computational discovery. Modern research workflows rely on the use of various evolving bioinformatics software applications which has led to a sophisticated computational ecosystem that accommodates increasingly diverse and integrated data sets along with their associated experimental probes, as well as comprehensive conceptual models of scientific understanding. As an example, consider an investigation of the mechanisms of promoting growth related to plant and microbe interactions as described in Finkel et al. [1]. They found a trait — such as the inhibition of root growth — can be significantly reversed by interaction with a single operon found in the bacteria *Variovorax*. The fact that *Variovorax* is involved is not surprising as it had previously been observed, but that such an important function is provided in a relatively simple way within a complex biological system, by a single operon, is unexpected. This investigation involved the use of multiple software tools to support data driven discovery and model based hypothesis generation [2].

A conceptual approach to reproducing their workflow might start with microbiome sequencing (performed in the wet lab), recovering metagenome assembled genomes or MAGs (using an assembly computational tool), annotation of the MAGs (using an annotation computational tool that performs alignment), followed by a taxonomic classification (using another computational tool), and followed by an analysis of associated biochemical pathways [3] (using a tool such as a flux balance analysis). Thus, this novel result required four different computational tools. As we demonstrate later, each of these tools may be considered almost as complex as the biological subsystems being explored.

The above example illustrates a single software workflow. Different research questions will require different workflows with different application parameters. These application *parameters* (or settings) can be coefficients or rate constants for biological processes, or they can be parameters to algorithms. This has led to the inclusion of many parameters in bioinformatics tools. We use the term *configuration* to refer to a specific set of chosen parameters. For instance, BLAST (*Basic Local Alignment Search Tool*) [4], one of the most commonly used tools to study DNA and protein sequences, has approximately 50 parameters a user can modify and customize. Changing these can impact core functionality such as increasing the string match length, or modifying the output format. Users can simply leave the default settings, or modify a few settings based on their laboratory norms, but this practice may result in the full power of these tools not being leveraged with less than optimal results or missed observations.

However, modifying parameters without a full understanding of their purpose can also be problematic. For example, in a letter to the editor of *Bioinformatics* by Shah et al. [5], they discuss a misunderstanding of a commonly used BLAST parameter (`max_target_seqs`) by another researcher. BLAST documentation states the following definition of `max_target_seqs` [6]:

> Number of aligned sequences to keep. Use with report formats that do not have separate definition line and alignment sections such as tabular (all outfmt > 4). Not compatible with num_descriptions or num_alignments. Ties are broken by order of sequences in the database.

Instead of this parameter filtering at the end as this may be commonly interpreted, it actually filters results during the algorithm’s iterative search. Thus it may not be clear to a user how it impacts their results. The authors of the referenced paper stated their explicit reason for limiting `max_target_seqs` to a value of 1, under the assumption it filtered out only the best (top) result at the end of the BLAST search. However, they misunderstood the parameter’s purpose. In a response from the NCBI team [7], they acknowledge that Shah discovered a bug (which they fixed) and that they have updated their documentation. Follow on work by González-Pech et al. [8] point to a number of misunderstood parameters of the tool.

In addition, several studies have examined how users interact with common bioinformatics tools. Morrison-Smith et al. noted that users are unsure of how to select the correct parameters and have to trust a tool’s correctness without real confidence [9]. For example participants have said “At the end of the day, you just have faith that the program’s working or not.” Others have examined how changing parameters impacts system performance [10] or correctness [11], however, there is a lack of research on how to train scientists to efficiently leverage the potential of parameters. We argue that if scientific software is viewed as a *magic box*, the community loses scientific potential. Instead, the bioscience user community needs to peel away the covers and develop techniques that explore the impact these parameters have on their results, managing them using a controlled and scientific approach. Reading documentation alone may not be sufficient.

Without a set of systematic techniques, it can be difficult to manage this software complexity. Since scientists use these tools to help form the basis of their conclusions, a lack of understanding of how parameters change the outcome of the results is both a missed opportunity and potentially problematic. In fact, it has already led to publication retractions [12, 13]. Biologists, hence, often develop a laboratory-specific routine protocol of what to change and leave the rest of the options alone. In this article we present a systematic and exploratory protocol (SOMATA) to work with scientific software. We also argue for more interaction between tool developers and end-users to build a shared body of knowledge. This will aid the advancement of computational methods to match the rigor that already exists in the documentation of lab work.

In the following sections, we first present an example and approach for working with bioinformatics software that matches the rigor and practice that already exists in the documentation and execution of laboratory work. Then we discuss background on software configurability and the bioinformatics tools used in this work. Next, we propose a systematic protocol (SOMATA) for breaking down the complexity of software tools. We then apply this protocol in a series of experimental studies with one study focused on empirically measuring the effect of using different configurations (set of choosen parameters when using a tool), and a second study demonstrating an approach to exploring the parameter space. Finally, we end with a discussion mapping out three key takeaways.

## Systems View - A Motivating Example^1^

A *systems view* in the biological context is a useful approach to study complex and potentially interacting systems. In this paper we take a systems view of software itself. Instead of using software as a single tool, we demonstrate how users can approach software with the same mindset as the organisms they study; as a complex system that can be optimized based on one’s environment and goals. We demonstrate this idea with an example showing how application parameters can change how a researcher perceives and interacts with bioinformatics tools while conducting a common experimental scenario. In this scenario we are trying to understand how different chemical compounds in a growth media change the metabolic pathways utilized in *Escherichia coli* (*E. coli*). For our growth medium we use the minimal media formulation Carbon-D-Glucose.

Before running extensive laboratory experiments we might want to exploit a common computational method simulating an organism’s growth using a genome-scale metabolic network [14]. We can run a Flux Balance Analysis (FBA) which calculates the flow of metabolites through the metabolic network in a specific growth media to optimize for growth (measured by flux through the biomass reaction) using a linear programming algorithm. The FBA tool used here (further discussed in experimental study methodology) provides input options such as the FBA model and the media, as well as algorithmic options such as the reaction to maximize, custom flux bounds, expression thresholds, and maximum uptakes of different nutrient sources. One output of this tool is the *objective value (OV)* which represents the maximum flow through the biomass reaction as a measurement of growth [15]. If we do not manually change any of the parameters (i.e. run the default configuration) *E. coli* produces an OV of 0.164692 mmol/g CDW hr.

For the purpose of this example we will explore the effect of the `max oxygen uptake` parameter which is described in the documentation as the “maximum number of moles of oxygen permitted for uptake (default uptake rates varies from 0 to 100 for all nutrients)”. This parameter has no fixed default value since the maximum is constrained by the media input. We will change this to a value of 0 to see what the behavior is. Under this new configuration, *E. coli* no longer grows (OV of 0 is returned). Next we increase this value in increments of 10 to observe how the OV changes. Results for values from 0 to 100 can be seen in Table 1. We see the OV increases linearly until it reaches the maximum (also the default growth) of 0.164692 when `max oxygen` is set to 40. If we set this value past 40, there is no added effect. We will explore why this is in a later study.

**Table 1.** Flux Balance Analysis (FBA) in *E. coli* in a Carbon-D-Glucose media. Objective Value (OV) is reported under various values of the max oxygen parameter.

| Parameter | OV |
| --- | --- |
| Default | 0.164692 |
| Max O 0 | 0 |
| Max O 10 | 0.053666 |
| Max O 20 | 0.107332 |
| Max O 30 | 0.160998 |
| Max O 40 | **0.164692** |
| Max O 50 | 0.164692 |
| Max O 60 | 0.164692 |
| Max O 70 | 0.164692 |
| Max O 80 | 0.164692 |
| Max O 90 | 0.164692 |
| Max O 100 | 0.164692 |

This example demonstrates changes in the measured result (OV), but there are also changes occurring to the internal model of *E. coli* — specifically — the flows or fluxes of each reaction in the metabolic network. The resulting flux can be positive meaning the net flow of metabolites occurs from left to right in the reaction equation, negative meaning the net flow occurs right to left, or zero indicating the flux is equal on both sides resulting in a net of zero. For example, reaction 1, below, shows the conversion of glucose to glucose-6-phosphate, a reaction with positive flux, from glycolysis.

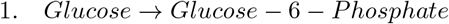

Reaction 2, below, shows this conversion in reverse as is the case in gluconeogenesis; this represents negative flux as glucose-6-phosphate is converted back to glucose.

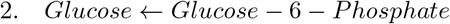

For more information on FBA within this tool please refer to [15]. The metabolic model used in this example has a total of 1,591 reactions. In modifying the level of oxygen, for example changing from the default to a value of 10, 431 reactions (27.09%) result in different fluxes.

Most reactions either stayed the same (72.91%), or increased or decreased in flux but their direction stayed the same (26.34%). However 12 reactions had a significant change as seen in Table 2. Five reactions change from a negative flux to a flux of zero, three from zero to negative, two zero to positive, and two changed from positive to zero. These 12 reactions represent significant changes that occur to metabolism of *E. coli* when oxygen levels are varied by means of the tool parameter.

**Table 2.** Change in individual reaction fluxes after setting the max oxygen parameter to a value of 10 versus the default of unset.

| KBase ID | KEGG ID/<br>EC Number | Effect on<br>Flux | Associated Pathways |
| --- | --- | --- | --- |
| rxn00515_c0 | R00722<br>2.7.4.6 | neg to zero | rn00230 Purine metabolism<br>rn01100 Metabolic pathways<br>rn01232 Nucleotide metabolism |
| rxn00616_c0 | R00848<br>1.1.99.5 | neg to zero | rn00564 Glycerophospholipid metabolism<br>rn01110 Biosynthesis of secondary metabolites |
| rxn00715_c0 | R00970<br>2.7.1.48 | neg to zero | rn00240 Pyrimidine metabolism |
| rxn00935_c0 | R01257<br>1.1.99.16 | neg to zero | rn00620 Pyruvate metabolism |
| rxn04792_c0 | R06983<br>1.1.1.284 | neg to zero | rn00680 Methane metabolism<br>rn01100 Metabolic pathways<br>rn01120 Microbial metabolism in diverse environments<br>rn01200 Carbon metabolism |
| rxn00611_c0 | R00842<br>1.1.1.8, 1.1.1.94, 1.1.1.261 | zero to neg | rn00564 Glycerophospholipid metabolism<br>rn01110 Biosynthesis of secondary metabolites |
| rxn01673_c0 | R02326<br>2.7.4.6 | zero to neg | rn00240 Pyrimidine metabolism<br>rn01100 Metabolic pathways<br>rn01232 Nucleotide metabolism |
| rxn09188_c0 | R10507<br>1.5.99.8 | zero to neg | rn00330 Arginine and proline metabolism<br>rn00332 Carbapenem biosynthesis<br>rn01100 Metabolic pathways<br>rn01110 Biosynthesis of secondary metabolites |
| rxn00931_c0 | R01251<br>1.5.1.2 | zero to pos | rn00330 Arginine and proline metabolism<br>rn01100 Metabolic pathways<br>rn01110 Biosynthesis of secondary metabolites<br>rn01230 Biosynthesis of amino acids |
| rxn08657_c0 | 1.1.3.15 | zero to pos | ec00630 Glyoxylate and dicarboxylate metabolism<br>ec01100 Metabolic pathways<br>ec01110 Biosynthesis of secondary metabolites<br>ec01120 Microbial metabolism in diverse environments |
| rxn04938_c0 | R07140<br>1.1.1.284 | pos to zero | rn00480 Glutathione metabolism<br>rn00680 Methane metabolism<br>rn01100 Metabolic pathways<br>rn01120 Microbial metabolism in diverse environments |
| rxn08656_c0 | 1.1.3.15 | pos to zero | ec00630 Glyoxylate and dicarboxylate metabolism<br>ec01100 Metabolic pathways<br>ec01110 Biosynthesis of secondary metabolites<br>ec01120 Microbial metabolism in diverse environments |

We can observe these changes to the reactions and corresponding pathways directly in the KEGG pathway maps as seen in Fig 1 which depicts Pyruvate Metabolism. The reaction through EC 1.1.5.4 changes from a negative net flux (left) to a net flux of zero (right). We also see reactions change in the magnitude of their flux. For example ECs 2.3.1.54, 2.7.1.40, and 2.7.2.1 change from a higher net negative flux (left) to a lower net flux (right). EC 2.3.1.8 changes from a higher net positive flux (left) to a lower net flux (right).

**Fig 1.**
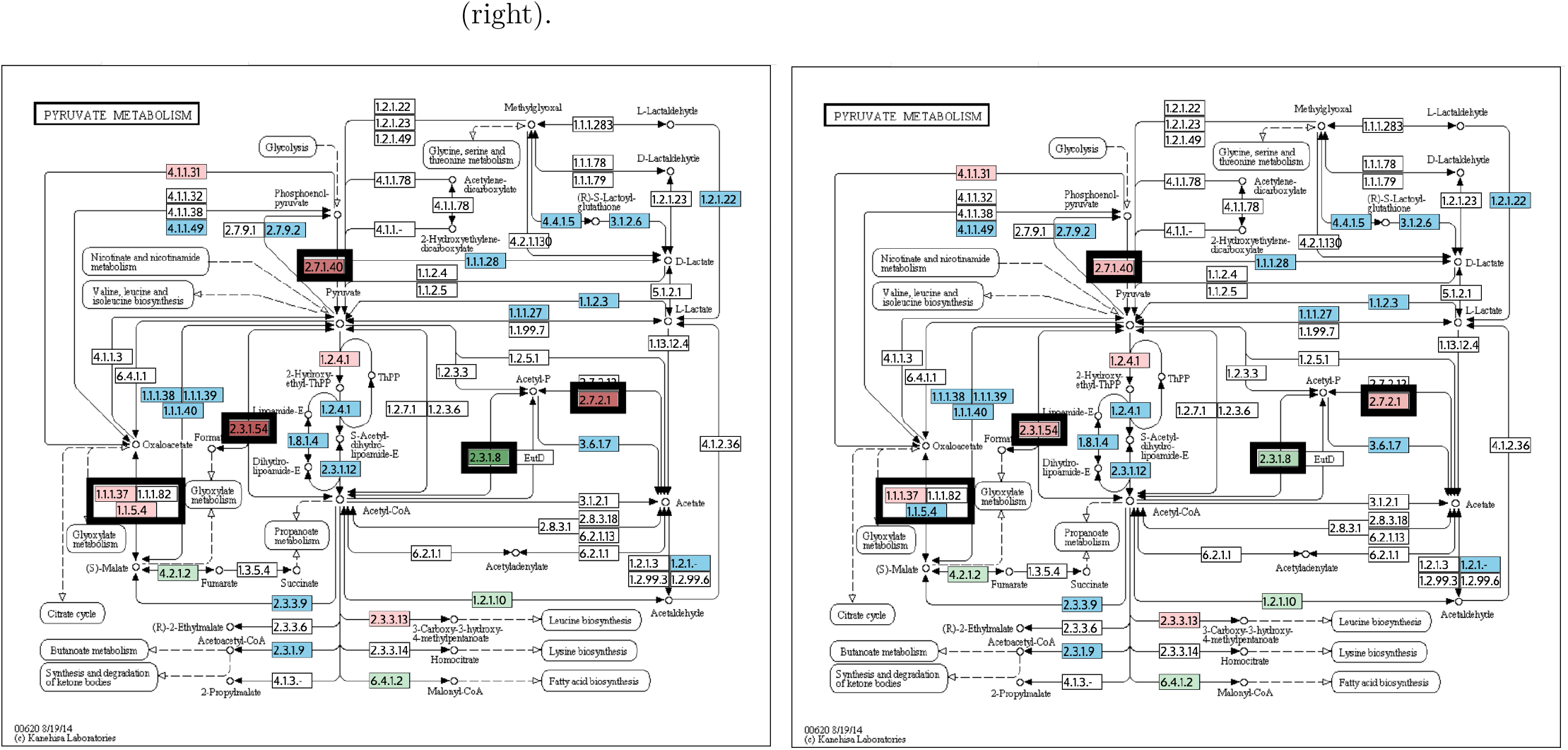
KEGG map for Pyruvate Metabolism under the default configuration (left) and with max oxygen set to 10 (right). Green represents a positive net flux, red is a negative net flux, and blue means the enzyme is present but the net flux is zero. The shade of the color represents the magnitude of the flux (a darker color represents a larger value).

These configuration changes can be considered as a “state change” to the system. Consider the state diagram in Fig 2. In the initial state (before running FBA) we have a static model with no defined fluxs through its reactions. After running the default configuration, we arrive at a state with a growth of 0.164692 with fluxes through its 1,591 reactions. But if we take a different transition, for example through `max oxygen` of 10 then we arrive at a different state with a growth of 0.0536659 where the flux through 431 reactions differ leading to changes in the pathways. The same occurs if we set `max oxygen` to 30 with a growth of 0.160998 resulting again in different reactions and paths. Hence, the configuration (and thus the parameters) have a direct impact on the individual reaction fluxes and the pathways through the network which leads to different behavior (e.g. growth).

**Fig 2.**
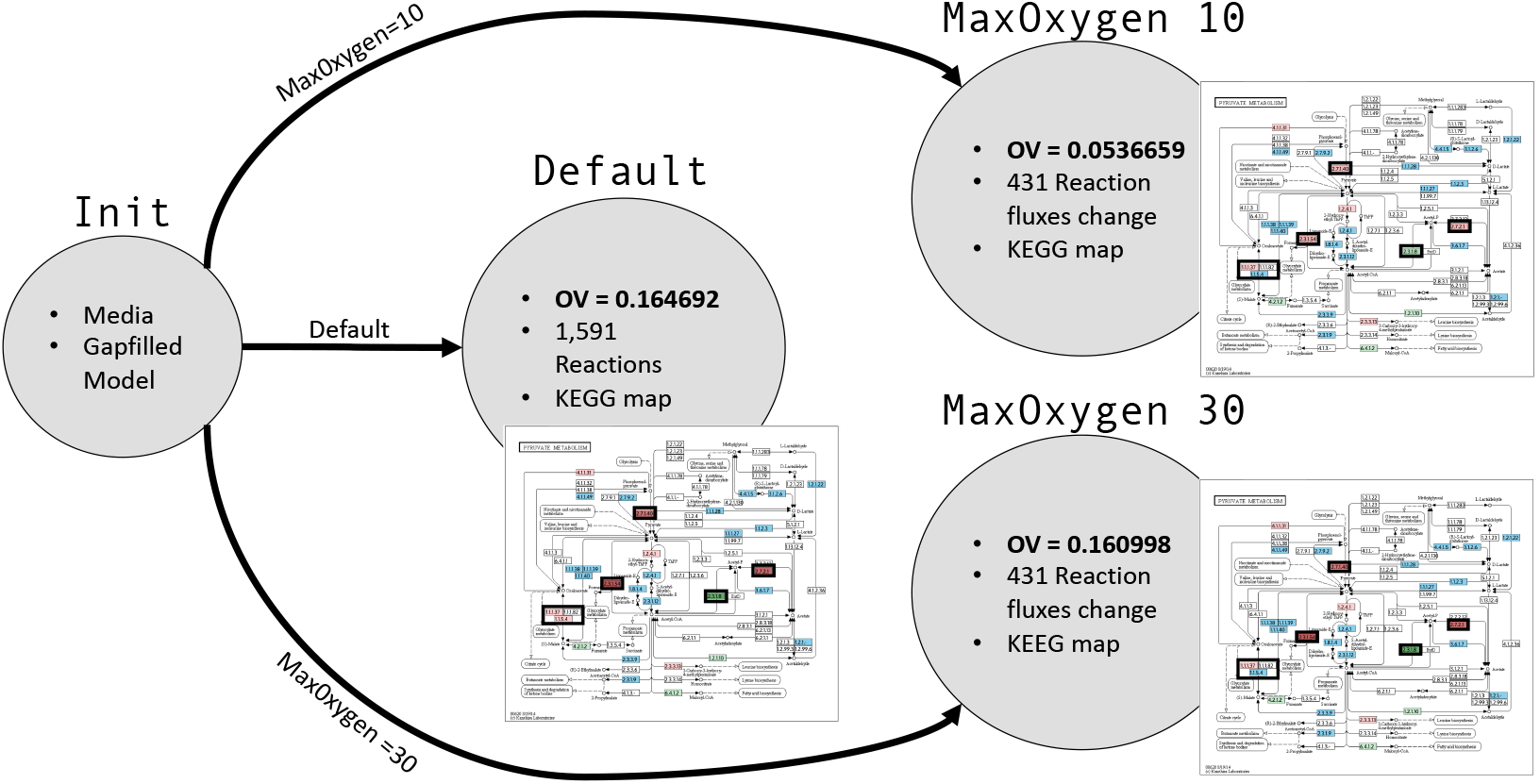
State change of FBA under different conditions.

By approaching this software tool with a systems view, we have demonstrated that not only can the ultimate objective change based on the parameters used (e.g. objective value), but the internal biological state (the direction and magnitude of the internal metabolic reactions) can change in significant ways as well. In the rest of this paper we demonstrate via case studies how using a systems mindset allows the scientist to explore and understand dependencies between application parameters and the final scientific result. This improved understanding can build confidence and lead to better science.

Next we present some background on configurability to lay the groundwork for our case studies.

## Background

### Software Configurability

Many software tools are designed with flexibility to satisfy a variety of use cases by supporting different facets of a user task. For instance, tools may handle different input data formats, various data sizes, and algorithms can be selected or customized depending on the user’s goal. A common example of configurability is a web browser, such as Firefox or Chrome. A user can select menu options that change the security settings or configure how windows and tabs open and close. Some users may want explicit warnings before exiting a window or tab, and others may prefer to close windows without confirmation. Which levels of security are chosen or the ability to embed JavaScript (or not) are all options that can be customized. Current versions of Firefox have more than 2,000 different options (e.g. parameters) a user can manipulate [16, 17].

Configurability benefits developers by allowing them to create software for a broad audience through building on reusable components, of which the cost of development can be amortized over the full software development lifecycle [18]. It also has been demonstrated to lead to higher quality of individual components since these are tested and used in multiple systems [18]. This type of development stands in contrast to building many unique specialized programs, each with individual code, and only slight variations in behaviors. From a user’s perspective, this would require them to switch between different software tools as their goals change.

A *configuration* of a software system is the set of choices, one for each of the system *parameters* (also known as *configuration options* or *settings*). For instance, in Firefox a configuration would be defined by the set of choices for all 2,000+ options. Most will have a default value that is assigned when the application starts, hence the user rarely changes more than a few of these on their own; in theory any combination of parameters can be used together. Hence, the complete *configuration space* of a configurable system is the set of all possible unique sets of parameter values (or the Cartesian product of the options for the individual values). In the simplest case, where all parameter are binary (turn the option on/off), the full configuration space would be 2|^*p*^| where |*p*| represents the number of parameters of the system. Returning to the Firefox example, assuming 2,000 parameters, this would be 2^2000^.

By design, changing parameters modifies how an application behaves. The possible space of behavior of a configurable software system is usually infeasible to exhaustively enumerate [11, 16, 20]. The range of system behaviors can lead to unexpected failures (both in outcome and performance) when interactions occur between parameters. For instance, one parameter might reduce the amount of memory for caching and another might increase some data that is being cached. When the new data is cached it might overflow the internal structures for caching which have now been reduced in size. There has been a large body of research that has identified efficient ways to sample a configurable software system’s configuration space and discover a large portion of the possible behavior [16, 21, 22]. We do not attempt to survey that literature here.

In this paper we focus on configurable software in the bioinformatics domain. We consider a *software parameter* (herein shortened to *parameter*) to be an option that can be specified by the user and then used by the program or application when it is running. As noted previously, users have highlighted difficulties understanding parameters, and this has even led to retracted articles [12, 13]. For a concrete biological example of a parameter, consider the graphical user interface of NCBI’s BLASTn implementation (Figure 3). This interface has more than 15 parameters for a user to adjust. In fact, BLASTn actually has over 50 parameters that can be modified thus not all are exposed in the NCBI GUI. Some of the parameters are in the form of a drop-down menu or a *set* of options such as `Max target sequences` and `gap costs`. Others are check boxes or *binary* parameters such as `short queries` and `Mask lower case letters`. Others are open text fields such as `Expect threshold` which can be set to a numeric value. Any parameter that is not explicitly defined will use the default value as determined by the program. While many laboratories have their own custom set of parameter option they regularly choose, there is no one best configuration that is best for all research questions, hence, understanding ways to navigate and explore these, is an important part of scientific discovery.

**Fig 3.**
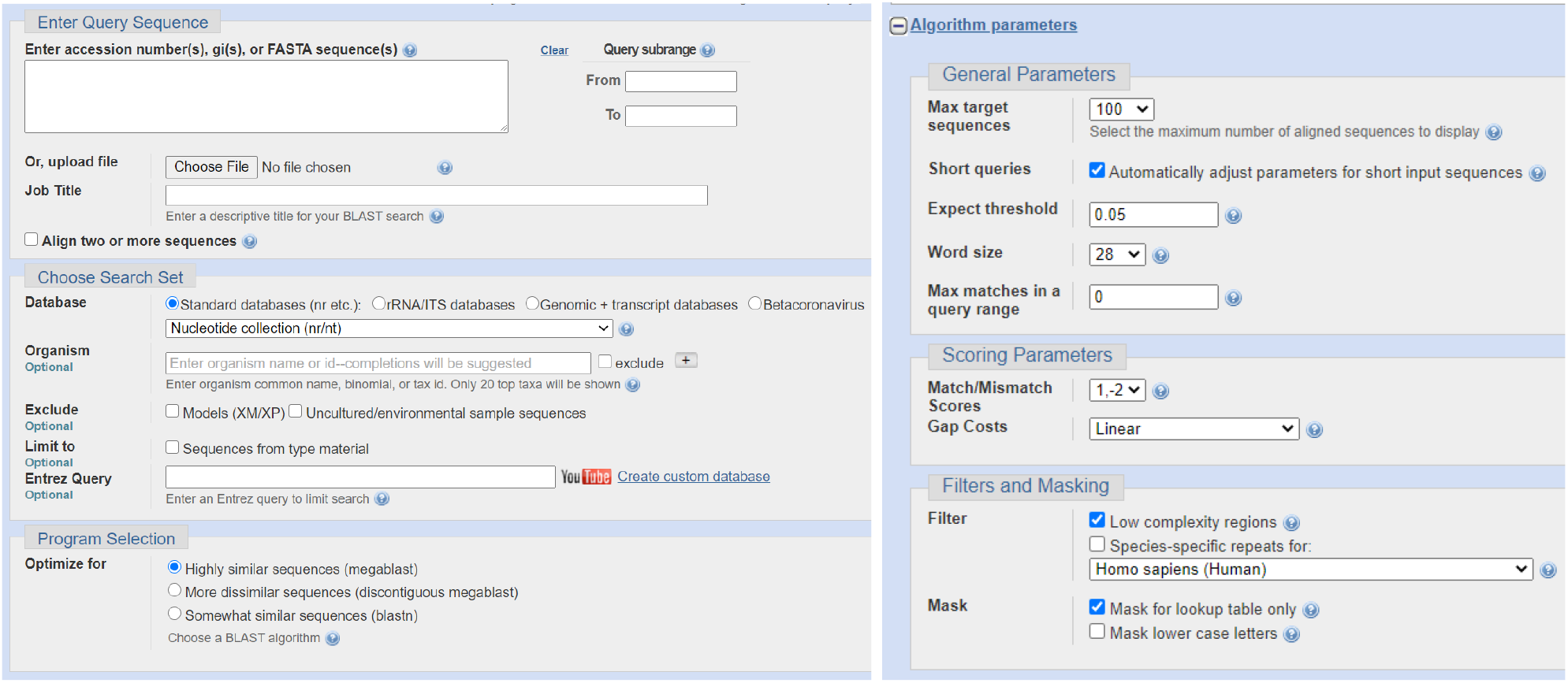
The Nucleotide BLAST (BLASTn) Graphical User Interface (GUI) environment from NCBI. The left shows the default input options, and the right shows the additional algorithmic parameters [19].

### Bioinformatics Methods

A common bioinformatics workflow, as shown in Figure 4, starts with *read files* output from a DNA sequencer that are then assembled (1) into reconstructions of the underlying original source genome DNA sequence. These are then aligned (2) against known DNA sequences to find regions of similarity with known genes and their functions. This provides a basis for identifying genes and annotation (3) of the original source genomes. The gene annotations then provide the basis for establishing genome-scale metabolic models (4) and doing flux balance analysis.

**Fig 4.**
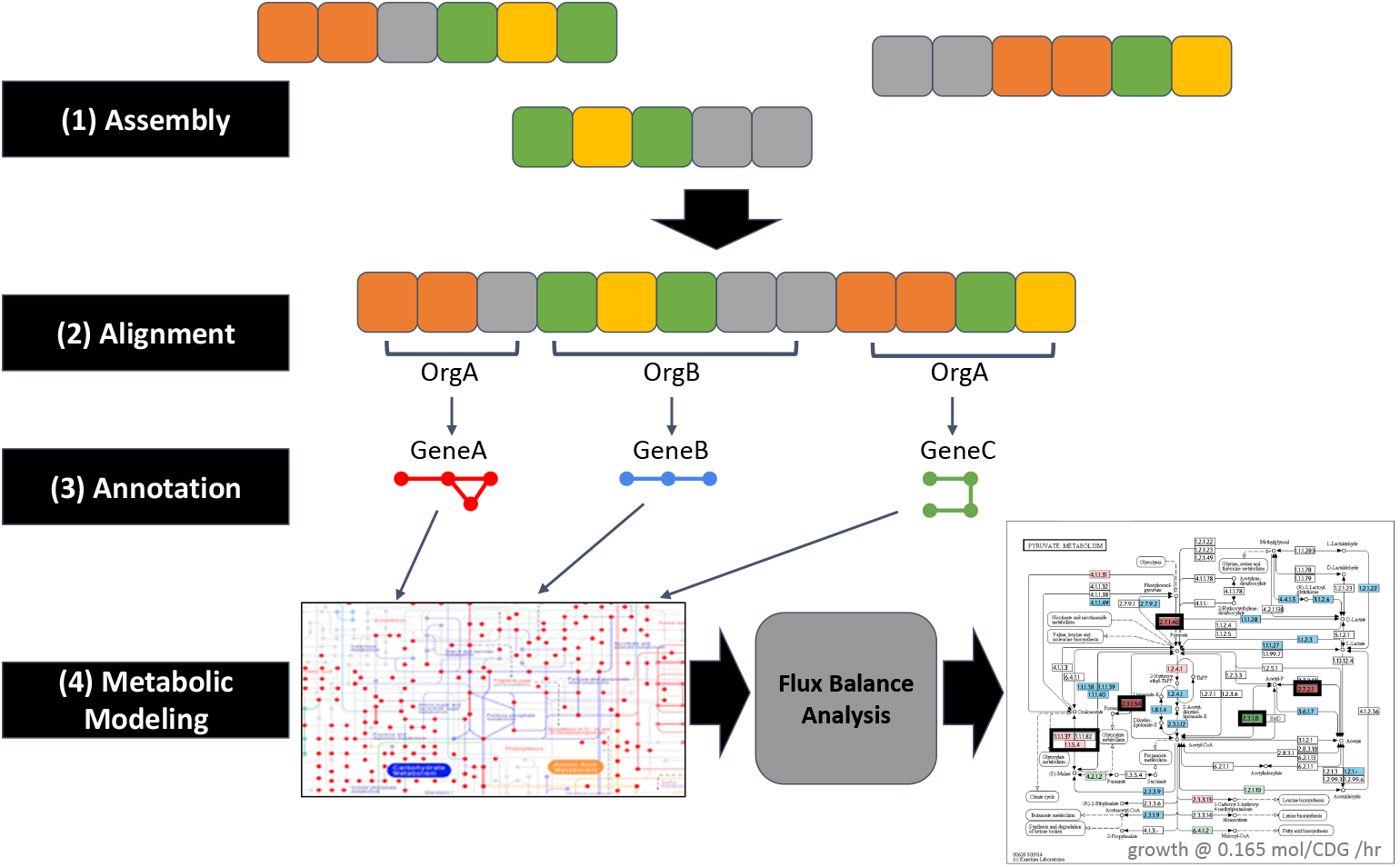
Methods overview of a common bioinformatics workflow.

In this work we consider three different exemplars of bioinformatics techniques commonly used in the workflow described: genome assembly, genome alignment, and metabolic modeling. The following subsections provide brief background on these techniques including a description of the biological functional objective of each. The term *functional objective* is used here referring to the biological output of the methods relative to the user’s biological query.

#### DNA Assembly

The *read file* output from a DNA sequencer contains a set of short sequences of base-pairs. Each of the short sequences is called a *read*, and are each an exact copy of a short segment of the DNA in the original sample. Depending on the sequencing technology and methods, each read ideally overlaps with many others. Assembly is the computational method to align and recombine reads to reconstruct the original genome sequence. When this is not fully accomplished due to sequencing error or lack of sufficient coverage, and especially with metagenomic samples containing genomes of many different species, the result is a set of assembled *contigs*, longer contiguous sequences reflecting the underlying genomes with hopefully high fidelity. A common **functional objective** of DNA assembly is to obtain a few long contigs, ideally a complete one, per species. For metagenomics there can be a trade off between completeness of individual species genomes and a more comprehensive representation of most species in the sample [23, 24]. Here we will focus on and optimize for the first.

#### DNA Alignment

Given a newly discovered or unknown DNA sequence, DNA Alignment (or Sequence Alignment) is a procedure for finding the most similar or homologous sequence in a database of known sequences [25]. The unknown sequence is often referred to as the query sequence. The alignment algorithm compares the query with all other sequences in the database, and calculates the significance of each match. We investigate a common use case, searching for the most similar known sequences in order to infer the likely evolutionary source or biological function of the unknown query sequence. The *matches* or *hits* found in the database to the query sequence are returned with a variety of quality scores that indicate how confident the alignment is with respect to each possible match. Two common metrics are the percentage identity (how identical two sequences are) and the expected value (or e-value) which is a measure of how likely the hits would be found in the database by chance. The **functional objective** of DNA alignment can vary by use case. We define one such objective that covers a range of use cases as the “number of quality hits” where the exact definition of quality may vary by use case, but typically involves a combination of several of the quality metrics such as percentage ID and e-value.

#### Metabolic Modeling

Metabolic modeling aims to reconstruct and simulate metabolic pathways of an organism to study its metabolism and gain further insight into its cellular biology. A metabolic network is a graphical representation of the set of chemical reactions where nodes represent chemical compounds and edges represent chemical reactions between those compounds. A path in these networks is a biological pathway. For example, the Kyoto Encyclopedia of Genes and Genomes (KEGG) database contains numerous metabolic networks for various model organisms [26]. One common type of analysis performed on metabolic networks is to estimate the flow of metabolites through the network under specific environmental conditions. A popular type of algorithm to perform this is known as flux balance analysis (FBA) [14]. FBA optimizes reaction fluxes (flows) through an organism’s metabolic reaction network to predict the organism’s growth or other objective function. As one example of a functional objective, a user might provide an FBA algorithm one environmental condition, and want to observe how much the organism grows. This can then be assessed by an *objective value* such as a measurement of biomass. Other examples of use cases include observing how pathways change under different environmental conditions, identifying what environmental conditions are needed for organism growth, and identifying how much an organism grows in a specific environment. The **functional objective** of FBA varies by use case. Herein we follow one of the most common; to determine the optimum flux over the metabolic network that maximizes growth.

## Experimental Protocol: SOMATA

We propose a process SOMATA, a core methodology to analyze any complex software systems and systematically break it down. SOMATA involves **S**electing tools and data, identifying **O**bjective metrics, **M**odeling the parameter space, choosing a sample design **A**pproach, **T**esting, and **A**nalyzing. SOMATA includes techniques from the field of software testing as its foundation. The details of each step are as follows:

1. **S**elect tool and data for exploration

- The first step is to select the tool or software of interest. In our motivating example the tool of interest was an implementation of Flux Balance Analysis, and the data was a metabolic model of *E. coli* along with a compatible definition of the Carbon-D-Glucose media.
2. Define an outcome metric based on user **O**bjective

- The metric to evaluate is an effect or output of the tool. The metric should be related to the user’s objective of interest. This could be a functional output such as a measurement associated with growth, as in our motivating example, or a non-functional output such as runtime. Several metrics may be selected depending on what is important in a particular use case. In our example the objective metric was the OV (a measurement of growth).
3. **M**odel the parameter space of the tool

- This involves identifying possible parameters of interest, defining their valid choices and range of options, and determining default values. Parameters can be environmental or functional parameters that have a known direct effect on the biology of the system, or they can be algorithmic parameters that are known to impact the underlying algorithmic method. In the example, a single biological parameter — `MaxO` — was varied since that is intuitively related to growth, the output objective of interest.
4. Choose sample design **A**pproach

- Next, choose from the constructed model in Step 3 which parameters to experimentally evaluate. A discrete range or set of options for each parameter must be chosen. Furthermore, the user must decide if they want to test the parameters individually, in combination, or some mixture of the two. In the example, we chose 11 choices for the parameter (values of `MaxO` between 0 and 100 in steps of 10) and explored these individually.
5. Execute the experimental **T**est evaluation

- Given the chosen model and sample design approach, run the selected configurations against the tool with respect to the selected data. Often this process can be automated to allow for large search spaces. Our example runs an FBA analysis for each of the 11 parameter choices.
6. Observe results and **A**nalyze effects

- Once complete, observe the effect of changing the model parameters given by Step 3, and conduct deeper analysis if desired. Given the results, to gain further insight, it may be beneficial to return to a previous step and alter the sections. For our example we examine the differences in OVs of growth and the impact on various associated pathways.

We want to emphasize the highly customizable nature of this process. For instance, in Step 4 there are several sampling design approaches that can be used. We can use techniques from software testing and design of experiments, or simply use a random method of investigation. In Step 5, simulation can replace experimentation. Finally, in Step 6 we could add in additional methods of analysis such as tools from machine learning. We follow this methodology in our Experimental Evaluation.

This process does not in itself contain any unprecedented elements individually. However as we argue in the Introduction, the bioinformatics and biology community would benefit from a clear approach to dissecting the complex software systems on which their research and science rely. Therefore, we present this as a formal methodology along with several experimental demonstrations of its application, and the real scientific impact that follow from the examples.

## Experimental Evaluation

In this section we expand on the scenarios in our motivating example, and ask how much of an impact changing parameters have on scientific results. For this we measure the behavioral impact of modifying parameters in several commonly used bioinformatics tools, which can lead to different scientific conclusions. We then ask if a systematic investigation of parameters can lead to any insights that can assist a user in navigating the parameter space. This would allow scientists to infer information to help them navigate the parameter space. Please refer to our supplementary website for study details and data^2^. Additionally, public KBase narratives demonstrating experimentation discussed here are provided^3^.

We formally ask the following research questions:

- **Experiment 1: What is the impact on a tool’s behavior when changing parameters?**
- **Experiment 2: Can we use an experimental approach to explore the parameter space and infer the internal behavior of the software system?**

We further split up Experiment 2 into two studies (Experiment 2a and Experiement 2b). We followup our study by sharing several lessons learned during this work that can lead to a set of best practices for users of highly configurable scientific software tools.

### Software Subjects (tools)

In Experiment 1 we leverage prior work which explored a set of four bioinformatics tools [27]. That work demonstrated the effect of parameters on various objectives, and identified software failures, but did not analyze the impact on the scientific outcome in depth. In this work, we provide an alternative analysis of the data and focus on the extent of the changes in the results, and whether they conform to scientific expectations.

We use an exemplar tool for each of the three bioinformatics applications described in the Background. For one of these applications we use two different versions of a tool resulting in four subjects in total. Criteria for choosing the tool subjects include: (1) the tool is frequently used in the community, (2) has a set of modifiable parameters, (3) can be run using automated scripts, and (4) includes a tutorial (or sample data) and documentation. We explicitly use data from others to avoid biasing our findings, and use tool documentation to guide parameter settings and ranges. For several of the subjects we utilize implementations from the U.S. Department of Energy’s System Biology Knowledgebase (KBase) which is a software and data science platform for researchers to analyze, integrate and model plant and microbial physiology and community dynamics [28].

The exemplar DNA alignment subject is the Nucleotide BLAST *Basic Local Alignment Search Tool* (BLASTn), a popular bioinformatics tool developed by the National Center for Biotechnology Information (NCBI) [4, 29]. We use the command line implementation of nucleotide BLAST (BLASTn), referred to simply as BLAST from here on [30]. The second subject is a DNA assembly tool. We chose the MEGAHIT algorithm [31, 32], and use a version implemented within KBase (version 2.2.8) [33].

The last two subjects perform Flux Balance Analysis (FBA). The first subject is the graphical user interface (GUI) app, *Run Flux Balance Analysis* [34], from within KBase’s main interface (herein referred to as FBA-GUI). It exposes only a limited set of all the possible parameters available in the underlying algorithm of FBA to be more tractable for new KBase users. Alternatively users may also adopt the command line interface (CLI) implementation of FBA (herein referred to as FBA-CLI). This FBA subject exposes many more parameters which can be manipulated. We utilize the open source standalone version (toolkit) of the FBA version 1.7.4 in this work [35].

## Experiment 1: Impact of Changing Configurations

This first study aims to answer the first research question: how does a tool’s behavior change when parameter settings are modified.

### Methodology

The methodology here follows the experimental approach previously outlined: SOMATA: **S**elect tool and data, choose **O**bjective metric, **M**odel parameter space, choose sample design **A**pproach, **T**est, and **A**nalyze. Starting with **step 1** we **S**elect the tools to be evaluated, we chose four subjects: BLAST, FBA-GUI, FBA-CLI, and MEGAHIT.

In **step 2** we define a set of functional outputs as **O**bjective metrics for each tool. For BLAST we use the number of *quality hits* using a strict definition of 100% identity value and and e-value of 0.0. This is a common measurement used to determine how well the query has performed. For FBA-GUI and FBA-CLI we use an objective metric that represents growth or biomass called the *objective value*. For MEGAHIT we selected four different metrics. The first two metrics count the *number of contigs* after a filter is applied. The metric *gt1kbp* is the number of contigs of length greater than one thousand base pairs, and the metric *gt10kbp* is the count greater than 10 thousand base pairs. Metric *N50* is the N50 score provided by MEGAHIT which is a weighted median statistic on the contig length. The last metric, *Max-bp*, is the length of the longest contig found. Note that these metrics can be adjusted to suit multiple use-cases. In this work we are simply restricting that choice.

Next in **step 3**, we define a **M**odel of the parameter space for each subject tool by selecting a sample of its parameters to test as seen in Table 3. For each tool we show the number of parameters, and for each parameter we list the type (i.e. Boolean, integer, float, or string) and state the number of choices used. For instance, we have two choices for Boolean parameters, three for strings and five each for integers or floats. As an example, if we examine BLAST we can see the model has three Boolean, three floats, and one string parameter for a total of seven parameters. This results in a combinatorial parameter space of 2^3^ × 5^3^ × 3 or 3, 000 different configurations. Please refer to our supplementary data for the exact parameters tested.

**Table 3.**
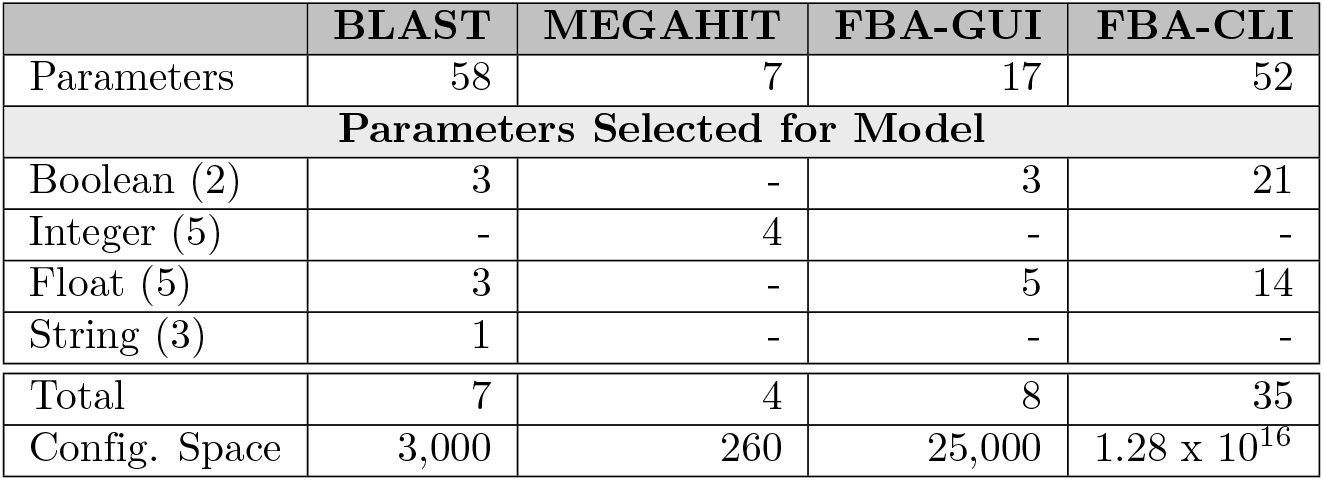
Subjects. For each subject we show the number of total parameters followed by the number chosen for testing broken down by parameter type. The number in parenthesis is the number of choices for that type. Any constraints on parameter values are not reflected here, please refer to our supplementary website for more information.

|  | **BLAST** | **MEGAHIT** | **FBA-GUI** | **FBA-CLI** |
| --- | --- | --- | --- | --- |
| Parameters | 58 | 7 | 17 | 52 |
| **Parameters Selected for Model** |  |  |  |  |
| Boolean (2) | 3 | - | 3 | 21 |
| Integer (5) | - | 4 | - | - |
| Float (5) | 3 | - | 5 | 14 |
| String (3) | 1 | - | - | - |
| Total | 7 | 4 | 8 | 35 |
| Config. Space | 3,000 | 260 | 25,000 | $1.28 \times 10^{16}$ |

In **step 4** we design an **A**pproach for sampling the parameter space. For BLAST, MEGAHIT, FBA-GUI we exhaustively test all combinations of their configuration options. For FBA-CLI the total configuration space is 1.28 × 10^16^ which is too many to exhaustively test. Thus we took a random sample of 125,000 configurations. Note that the configuration space for MEGAHIT of size 260 has been reduced from the full space of 625 due to constraints between parameters. Please refer to our supplementary data for more information.

To measure the impact of changing parameters we measure several data points to observe the number of configurations that change the output (for better or for worse). We can also look at the entire configuration space given our input data and research objective, and ask how hard it is to find a better, or even the best result compared to the default value.

### Results

After executing the experiment (**T**est in **step 5**), Table 4 presents the results for our **A**nalysis of the effects in **step 6**. Each row represents one subject tool and objective metric combination. Table 4a reports the number of configurations tested, the number of unique output metrics, and a summary of the range of output metrics (default, best, worst, and most common). This demonstrates the impact on the raw objective metric demonstrating the effect the configuration space has on the scientific output. Table 4b breaks down the percentage variation to give an idea of the landscape of the outputs.

**Table 4.** Analysis of objective metric impacts. **(a)** Objective Metric Impacts. *Model* is the tool (and objective metric) used. The *No. Configs Tested* column shows the number of configurations tested. The *No. Unique* column is the number of unique output values from all configurations tested as an indication of the diversity of output. The next four columns are the raw objective metric output values for the default, best, worst, and most common output values. **(b)** Percentage Variation Summary. *Model* is the tool (and objective metric) used. The *No. Configs Tested* column shows the number of configurations tested. The remaining columns represent the percentage of configurations tested with respect to the default value. For example *Better* represents the percentage of configurations that returned an objective metric better than the default. The final column is the percentage of configurations that resulted in an error by the software.

| Model | No. Configs Tested | No. Unique | Objective Metric |  |  |  |
| --- | --- | --- | --- | --- | --- | --- |
|  |  |  | Default | Best | Worst | Most Common |
| BLAST<br>(no. quality hits) | 3,000 | 17 | 2,569 | 4,351 | 1,809 | 2,569 |
| FBA-GUI<br>(OV) | 25,000 | 4 | 0.508 | 0.508 | 0 | 0 |
| FBA-CLI<br>(OV) | 125,000 | 39 | 0.508 | 1.171 | 0 | 0 |
| MEGAHIT<br>(gt1kbp) | 260 | 122 | 284 | 656 | 2 | 0 |
| MEGAHIT<br>(gt10kbp) | 260 | 47 | 143 | 147 | 0 | 0 |
| MEGAHIT<br>(N50) | 260 | 147 | 17,227 | 17,593 | 0 | 0 |
| MEGAHIT<br>(Max-bp) | 260 | 88 | 64,476 | 70,112 | 0 | 0 |

| Model | No. Configs Tested | Default | Better | Best | Worse | Worst | Most Common | Errors |
| --- | --- | --- | --- | --- | --- | --- | --- | --- |
| BLAST<br>(no. quality hits) | 3,000 | 17.33% | 72.00% | 13.33% | 10.67% | 2.00% | 17.33% | 0.00% |
| FBA-GUI<br>(OV) | 25,000 | 18.43% | 0.00% | 18.43% | 81.57% | 67.23% | 67.23% | 0.06% |
| FBA-CLI<br>(OV) | 125,000 | 0.56% | 1.58% | 0.05% | 97.60% | 95.14% | 95.14% | 1.03% |
| MEGAHIT<br>(gt1kbp) | 260 | 1.15% | 66.15% | 0.38% | 9.62% | 0.38% | 23.08% | 0.00% |
| MEGAHIT<br>(gt10kbp) | 260 | 8.46% | 17.69% | 0.38% | 50.77% | 7.31% | 30.38% | 0.00% |
| MEGAHIT<br>(N50) | 260 | 2.31% | 2.69% | 0.38% | 95.00% | 0.38% | 23.08% | 0.00% |
| MEGAHIT<br>(Max-bp) | 260 | 0.38% | 13.46% | 1.54% | 63.08% | 0.38% | 23.08% | 0.00% |

In Table 4a we can see the number of unique objective values is low compared with the total number of configurations tested in most subjects, suggesting random sampling may be very helpful. However in MEGAHIT, for the N50 and gt1kbp metrics, we see about half of the configurations leading to unique objective values. This may suggest we need to create larger samples to fully explore all potential behaviors of those objectives.

Looking at the range of the objective metrics (last four columns), we can see in some cases there is a large increase in the result comparing the default output to the best (BLAST, FBA-CLI, MEGAHIT gt1kbp). Other subjects show a smaller range. However all subjects have a substantially lower worst metric, and all except BLAST have an output of zero as their most common output metric.

In Table 4b we can see the landscape of the distribution of the scores varies greatly on the subject and objective. For BLAST 72% of the configurations improve over the default, but the best result is only seen in 13.33% of the configurations. In FBA-GUI the default is the best value, but that is only achievable by 18.43% of configurations. In FBA-CLI - the largest configuration space - only 1.58% of the random configurations improve over the default’s output, and 0.05% result in the best output value. In fact most of the time (in 95.14% of configurations) the worst value (0 or no growth) is achieved. For MEGAHIT the odds of improving over the default range from 2.69% (for N50 score) to 66.15% for the number of contigs greater than 1,000 base-pairs. Finding the optimal value for any MEGAHIT metric is low across all subjects ranging from a 0.38% to 1.54%. We see that the number of configurations that achieve the same result as the default ranges across subjects, as low as 0.38% for MEGAHIT (measured by maximum base pairs) and as high as 18.43% for FBA-CLI. In all subjects more than 80% of the results are different from the default.

#### Summary

These results show that the default configuration does not always provide the best objective value, and that instead of randomly choosing configurations, a more systematic method of investigation is needed. For example in all subjects except BLAST, if a user randomly chooses a configuration, most often the worst value (0) will be obtained. **The variation in the value of the result also demonstrates the importance of reporting the exact configuration used in any documentation or publications; otherwise the results are not reproducible.** The ranges in the configuration distributions in MEGAHIT tell us several things. We can see that the odds of improving over the default N50 score is significantly less than the other use cases. This signals that MEGAHIT may be optimized for the N50 score over other use cases. However, if a user is more concerned with achieving more contigs with sizes greater than 1,000 base-pairs they may need to consider other configurations. Configurations are not one-size-fits-all. This demonstrates how essential it is for the user to understand their particular use case, have an effective metric that is representative of the objective value, and consider how different parameters may affect it.

## Experiment 2: Experimental exploration approach

Motivated by the results of Experiment 1 demonstrating tool configurations can have a large impact on the scientific conclusion, in this next study we ask how a user might use an experimental approach to explore the parameter space and infer the internal behavior of the system. We do so through two studies. In Experiment 2a, we focus on a single parameter to explore in depth how this parameter affects an objective. In the process we uncover some unexpected behavior. In Experiment 2b we explore the overlap in functionality between multiple parameters. We noticed that some parameters can be changed from either setting a single parameter within the GUI interface, or alternative sources. We wanted to understand the interplay between these choices. Experiment 2a demonstrates how a user can systematically probe a software tool to verify and learn the impact of single parameters. Experiment 2b provides a method to explore the interaction between multiple parameters.

### Experiment 2a: In-Depth Study of Single Parameter

We extend the experiment from our motivating example on two additional subjects to observe if the effect is the same or different.

#### Methodology

Following **step 1** of our protocol we **S**elect the KBase GUI implementation of FBA as the subject tool and select three subject organisms that come from tutorial or public data in KBase. We tested both *Escherichia coli* (*E. coli*) and *Rhodobacter sphaeroides* (*R. sphaeroides*) growing in the standard Carbon-D-Glucose media. We also tested *Shewanella oneidensis* (*S. oneidensis*) in the Lactate media. Please refer to our supplementary material for an in-depth description of the experimental design. Per **step 2**, we use the same **O**bjective metric to evaluate as Experiment 1, growth measured by the objective value, in all cases. For our **M**odel of the parameter space (**step 3**), we investigate the use of a single parameter the `Max Oxygen` parameter (maximum number of moles of oxygen permitted for uptake) which according to the specifications can be set between 0 and 100. The default has no explicit set value. We determine the sample design **A**pproach (**step 4**) by systematically testing a range of values (from 0 to 100 in steps of 10) for `max oxygen`. In order to test the bounds of the parameter, we also tested values larger than the maximum according to the documentation (100), thus we also tested values of 101, 250, 500, and 1000.

We consider a use case where the user may want to identify what setting for the max flux of oxygen results is the highest growth. However they do not want to oversupply the levels of oxygen, so they want to identify the minimal value of the flux of oxygen to achieve the highest OV.

#### Results

After executing the experimental **T**est in **step 5**, Table 5 presents our results for all three organisms for **A**nalysis in **step 6**. *E. coli* and *R. sphaeroides* both showed similar behavior. The growth for each starts at 0 when `max oxygen` is set to zero, then increases to the default growth at a `max oxygen` value of 40 for *E. coli* and 50 for *R. sphaeroides*. Any value higher results in no additional growth.

**Table 5.** FBA varying MaxO

| Configuration | E. coli (CDG) | S. oneidensis (Lactate) | R. sphaeroides (CDG) |
| --- | --- | --- | --- |
| Default | 0.164692 | 3.271000 | 0.545390 |
| Max O 0 | 0 | 0 | 0 |
| Max O 10 | 0.053666 | 0.113570 | 0.120263 |
| Max O 20 | 0.107332 | 0.227139 | 0.240526 |
| Max O 30 | 0.160998 | 0.340709 | 0.360790 |
| Max O 40 | **0.164692** | 0.454278 | 0.481053 |
| Max O 50 | 0.164692 | 0.567848 | **0.545390** |
| Max O 60 | 0.164692 | 0.681418 | 0.545390 |
| Max O 70 | 0.164692 | 0.794987 | 0.545390 |
| Max O 80 | 0.164692 | 0.908557 | 0.545390 |
| Max O 90 | 0.164692 | 1.007780 | 0.545390 |
| Max O 100 | 0.164692 | 1.099590 | 0.545390 |
| Max O 101 | 0.164692 | 1.108690 | 0.545390 |
| Max O 250 | 0.164692 | 2.123170 | 0.545390 |
| Max O 500 | 0.164692 | **3.271000** | 0.545390 |
| Max O 1000 | 0.164692 | 3.271000 | 0.545390 |

*S. oneidensis* also grows at a linear rate starting from 0 growth at a `max oxygen` value of 0. However it does not reach the default growth at the maximum value of 100. However, if we go beyond the documentation specified range, the default growth can be achieved by setting `max oxygen` to 500. Surprised by this behavior we asked the developers of this tool for further information where we learned that the documentation was misleading and that 100 is not a hard maximum value.

##### Summary

Using a systematic approach we were able to verify the expected behavior that increasing the maximum allowed oxygen flux would allow for an increase in organism growth. **This increases the user trust in the tool and verifies our understanding is in line with the tool’s functionality.** We were also able to identify the smallest value of this setting that results in the best scientific output (40 for *E. coli*, 50 *R. sphaeroides*, and 500 for *S. oneidensis*). This answers our use case question. Furthermore, we identified some unexpected behavior that was not in line with the documentation. **Tools are not without bugs or mistakes in documentation and it is useful for users to be equipped with alternative methods to be able to identify any unexpected behaviors on their own data and use case, and clarify them with developers.**

### Experiment 2b: Interaction between parameters and the environment

As previously noted, we identified the flux limits can be altered in more than one manner. In this experiment we further investigate how these different methods interplay. One way to control flux is within the media input file via a field for *max uptake* of each compound with units “mol/per CDW hr”. As a second method, from the GUI the carbon fluxes can be constrained via an application parameter. There are five key nutrients that can be restricted: carbon (C), nitrogen (N), phosphate (P), sulfur (S), and oxygen (O). We will refer to these as `MaxC`, `MaxN`, `MaxP`, `MaxS`, and `MaxO` respectively. The description of these parameters is “Maximum number of moles of [C/N/P/S/O] permitted for uptake (default uptake rates varies from 0 to 100 for all nutrients)”. There is yet a third way to constrain the fluxes. We can also set individual limits on either compounds or reactions using a different (string) parameter (`Custom Flux Bounds`). We will explore the first two methods. The third method can be explored using a similar approach.

This experiment investigates how changing different parameters affects the final scientific result. For example if there is a limit of 10 on MaxO through the application parameter, and a limit of 40 for *H*_2_*O* specified by the media, what is the resulting limit for the exchange flux of *H*_2_*O*? Does one supersede another? Or is the minimum or maximum taken as the limit? How is the flux of multiple compounds affected if they all share common nutrients? The impact of such changes may be non-trivial to understand without a systematic exploration. Furthermore, the output of the FBA tool produces multiple values for exchange fluxes including *upper bound*, *max exchange flux*, and *exchange flux*. It may not be obvious to a user what precisely each of these fields mean and where the limits are coming from. Two specific questions we investigate in this study include:

1. If the *max flux* GUI parameters are left as unset, what source does FBA use to determine the limits on the uptake of the different carbon?
2. How can we identify the minimal, required uptake rate for each nutrient?

#### Methodology

Following **step 1** we **S**elect as the subject tool Flux Balance Analysis with *E.coli* in the Carbon-D-Glucose media as the base media. The outcome metric (**step 2**) remains as the **O**bjective value. In order to explore the sources of the flux constraints, the first step was to identify what compounds in the media the important nutrients could be coming from. For carbon, nitrogen, phosphate, and sulfur there is only one source for each: D-Glucose (C6H12O6), NH3 (H4N), Phosphate (HO4P), and Sulfate (O4S) respectively. Oxygen has several sources: Dioxide (O2), Sulfate (O4S), Phosphate (HO4P), H2O, Molybdate (H2MoO4), D-Glucose (C6H12O6). For simplicity, we chose two cases where there was only one source: carbon (MaxC) with source D-Glucose and nitrogen (MaxN) with source NH3. We left the media limits fixed (5 for D-Glucose and 100 for NH3), and then varied the max flux limits of carbon and nitrogen respectively. This defines our **M**odel of the parameter space (**step 3**).

For the sample design **A**pproach in **step 4** we take a different approach than the previous studies. We take a more iterative exploratory approach by starting with the default settings (unset in this case), altering them by a value of 10, and reassessing based on the results. In this type of a design, we started with an initial model (default as well as increments of 10), experimentally **T**ested (**step 5**), then observed and **A**nalysed (**step 6**) and went back to add to the **M**odel (**step 3**) and re-evaluate. Since there was no change as the Max uptake of N was increased from 10 to 20 and the max exchange flux was observed to be 1.5551, further exploration was performed by limiting the max uptake of N at values between 1 and 2.

#### Results

Results for exploring the `MaxC` parameter can be seen in Table 6. The rows *upper bound*, *max exchange flux*, *exchange flux*, and *OV* are all output values of the FBA object. Based on these results we can extrapolate about how the tool is working.

**Table 6.**
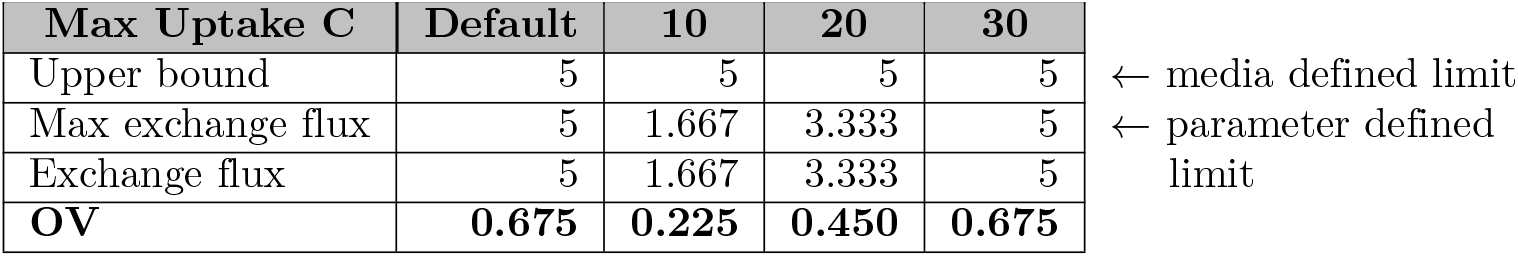
Max Uptake C Exploration - D-Glucose (C6H12O6)

| Max Uptake C | Default | 10 | 20 | 30 |  |
| --- | --- | --- | --- | --- | --- |
| Upper bound | 5 | 5 | 5 | 5 | ← media defined limit |
| Max exchange flux | 5 | 1.667 | 3.333 | 5 | ← parameter defined limit |
| Exchange flux | 5 | 1.667 | 3.333 | 5 |  |
| OV | 0.675 | 0.225 | 0.450 | 0.675 |  |

We can see that the *upper bound* is consistently at a value of 5, this must mean the upper bound is set by the media file. The field for *max exchange flux* however varies. Under the default parameter we see it is identical to the *upperbound*. However when we constraint the amount of `MaxC` this value varies. It is reduced in `MaxC`=10 and `MaxC`=20, but at `MaxC`=30 we see it is again equal to the default of 5. When *E.coli* has at least 5 units of carbon, it will grow at is maximum amount (0.675). Furthermore, we can see both the *uptake* and the OV increase linearly. At a `MaxC` of 10 the uptake is 1/3rd of the uptake under the default, and the OV is also 1/3rd of the OV under the default. At `MaxC` of 20 the uptake and OV are 2/3rds of the corresponding values under the default.

We repeat this process for nitrogen seen in Table 7. In this case we vary the nitrogen supplying compound (NH3) initially at levels of 10 and 20. In follow-up exploration, we also used levels of 1, 1.5, 1.5551, and 2. At 2 units and above, *E.coli* grows at is maximum (default) amount.

**Table 7.** Max Uptake N Exploration - NH3 (H4N)

| Max Uptake N | Default | 1 | 1.5 | 1.5551 | 2 | 10 | 20 |
| --- | --- | --- | --- | --- | --- | --- | --- |
| Upper bound | 100 | 100 | 100 | 100 | 100 | 100 | 100 |
| Max Exchange flux | 12.9392 | 1 | 1.5 | 1.5551 | 2 | 10 | 20 |
| Exchange flux | 1.55528 | 1 | 1.5 | 1.5551 | 1.55528 | 1.55528 | 1.55528 |
| OV | 0.164692 | 0.116501 | 0.164269 | 0.16469 | 0.164692 | 0.164692 | 0.164692 |

We observe that the exchange flux for NH3 is limited to 1.55528 as this is the amount of uptake for 2 through 20 units of nitrogen as well as the default case. Further, *E.coli* grows at its maximum (default) amount for these amounts of NH3. If we set a value slightly smaller than that as our `MaxN` (e.g. 1.5551) we see that it now grows at just 0.000002 less than its maximum potential. Further, at `MaxN`=1 it grows at 70.74% of its maximum amount while at `MaxN`=1.5 it grows at 99.74% of its maximal potential.

##### Summary

This experiment reverse engineered the effect of changing max flux parameters in FBA, and investigated how the application parameter interacts with the media options. **We were able to extract the meaning of two of the output values that were unclear from documentation.** We found that *upper bound* is the flux bound by the media and *max exchange flux* is bound by the application parameter. We found that the *max exchange flux* is correlated with which compounds in the media contain those nutrients, and that the minimum value between the two is used as the limiting value. We can also identify the minimal required exchange flux by looking at the actual exchange flux reported by the output. **This process shows how a user can systematically perturb a scientific software system in order to reverse engineer pieces of the underlying algorithm.** It was initially unclear from documentation how the interaction between these options operated, and what the meaning of the different output fields meant. By using this experimental approach the user was able to discover the answers on their own. Similar experimental designs could be set up to ask different types of questions.

## Discussion

We have seen using three exemplar experiments that biologists can benefit from approaching their computational tools the same way they approach their bench experiments. Instead of using tools as magic boxes, we propose SOMATA, an exploratory approach that leads to computational understanding and expertise. By leveraging application parameters (which were developed for specific use cases), a biologist can better understand the range of a tool’s application and thereby optimize the results for their use case. However, this is also a cautionary tale. Modifying the configuration of a tool may change how the tool works in unintended ways, and doing so in an ad-hoc manner can lead to results that are not reproducible or are potentially sub-optimal.

We also demonstrated that some parameters, while well documented, provide inaccurate information. While the tool authors document the valid values, these were in fact incorrect limits for the tool. We discovered via experimentation, and confirmed with the developers, that we can use larger values in the application, and that larger values result in (correct) larger values of growth. Situations like this can lead to a confusing landscape for biologists. Hence, we suggest using a FAIR-like (Findable, Accessible, Interoperable and Reusable) approach [36] not only with data collection but also to using and reporting interactions and final settings with all tools, just as FAIR has been proposed to be extended to research tools [37].

The FAIR initiative came about in recognition that science was being impeded by the lack of reproducibility caused by lax data standards. In a similar way, the advancement of research may also be hampered by ad hoc use of software tools that are increasingly becoming an integral part of scientific investigation. For instance Shade and Teal proposed a generic method for approaching workflows which could be considered a high-level abstraction of SOMATA [38]. As noted in the original article on FAIR [36] the principles should be applied equally to workflows and software tools, and data: “All scholarly digital research objects [39] — from data to analytical pipelines—benefit from application of these principles, since all components of the research process must be available to ensure transparency, reusability and reproducibility.” Therefore, it is incumbent on researchers to have a working understanding of the tools used in the context of their research, and to properly investigate and communicate their findings in a manner adhering to the principles of the scientific method.

We now synthesize and discuss some of the takeaways (or key lessons learned) during this study.

1. **Importance of Application Parameters.** The key takeaway from this study is that application parameters can have a direct impact on the output and can effectively change the biological *state* of the experiment. Thus, it is essential a user understand the impact each parameter has. Furthermore, if a user does not take advantage of the variety of parameter options, they may be missing out on functionality. On the flip side, we have also seen that having a limited understanding (or simplified/incorrect documentation) of different parameters may cause problems, hence the burden should not be solely on the biologist. Configurations matter, and the creation, documentation, and experimentation with them should be a community effort. We envision community resources where information from both experimentalists and tool developers interact. Rather than simply FAIR we suggest we also need to adopt a SHARE (Simple, Helpful, Accessible, Readable with Explanations) principle. First, we need methods that are easy to implement and understand so that biologists use them. Second, our systems should provide more hints (or help) to aid users in understanding the various configuration options. Third, the differences (or impact of) changing application parameters should be annotated with readable explanations. None of this can be achieved in isolation, hence we think this is a community effort.
2. **Mapping Configurations to Behavior May not be Obvious.** If we return to the documented `max_target_sequence` parameter in BLAST (discussed in the Introduction), we saw that a user could easily interpret the use of setting that option to represent an output filter after the search returns some number of hits; hence setting it to a small number could provide just the best results. But in fact, this parameter is used internally to filter results early in the algorithm, hence it may cause the search to perform sub-optimally. Why this parameter exists is out of the scope of this paper, but it was likely added to improve performance (i.e. make the search run faster). When that objective is required, then a less than optimal search may be beneficial. Had the author of that paper used our approach to explore parameter options they would have seen inconsistent results when setting this parameter (in fact it would only ‘by chance’ return only a single hit). The lesson here is that relying on simplified (or only) documentation could be deceptive.
3. **We Need Better Tools.** While we propose the use of exploration in this paper, we also see the need for next generation tools which provide explanations and expose configuration behavior. As science continues to advance, computational tools are increasingly vital to discovery. In order to maximize their benefit, we should strive to ensure they perform in an explainable, transparent, and intuitive way.

## Conclusion

In biological research, as the variety and resolution of data grows to increasingly reflect the complexity of biology, so too are the associated computational tools becoming more comprehensive mirrors of the same complexity. In order to investigate the increasing biological complexity, laboratory protocols are shifting not just through reductionism, but also taking an increasingly more comprehensive systems approach. In a similar approach here, we propose through SOMATA taking a systems view to bioinformatics research and tool development that mimics how biologists explore organisms in the laboratory. This provides a common basis for communication and interaction that will help the research community more productively develop and use those tools, and thereby advance their research efforts.

We have demonstrated that different results can be obtained using different algorithm and tool parameters and that this is a potential minefield for the novice user. Rather than limit exploration, we argue that the community of bioinformatics researchers should provide open, explainable tools and that this information should be shared and crowd sourced to provide a more repeatable environment. While the FAIR principles are of value here, we see a need for a more comprehensive way to view configurability in software tools.

A systems approach to biology implies the need for this same kind of interchange within the research community especially as the computational tools are refined to be more precise reflections of biological elements and the experimental processes investigating them. Having a common framework for decomposing these increasingly complex tools that mimics a systems view of biology is the kind of standard that we believe would substantially advance the overall scientific research effort toward the future.

## Supporting information

**Table 8.**
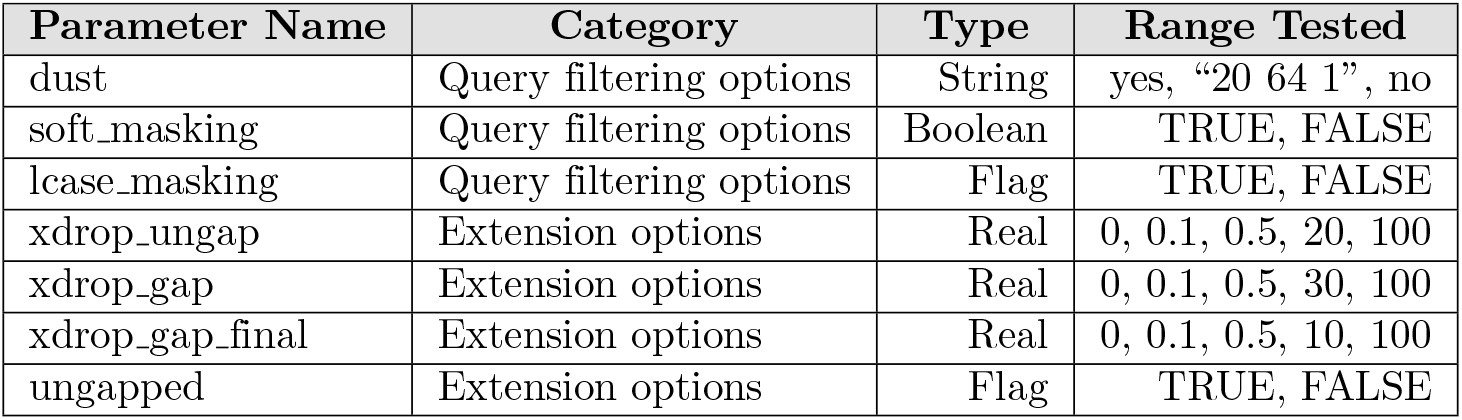
BLAST configuration model

**Table 9.** MEGAHIT configuration model

| Parameter Name | Type | Range Allowed | Range Tested |
| --- | --- | --- | --- |
| min-count | Numeric | [2,10] | 2, 4, 6, 8, 10 |
| k-min | Numeric | odd in range [1,127] | 15, 21, 59, 99, 141 |
| k-max | Numeric | odd in range [1,255] | 15, 21, 59, 99, 141 |
| k-step | Numeric | even in range [2,28] | 2, 6, 12, 20, 28 |

**Table 10.** FBA-GUI configuration model

| Parameter Name | Type | Range Tested |
| --- | --- | --- |
| flux variability analysis | Boolean | TRUE, FALSE |
| minimize flux | Boolean | TRUE, FALSE |
| simulate all single KOs | Boolean | TRUE, FALSE |
| max Carbon uptake | Float | 0, 25, 50, 75, 100 |
| max Nitrogen uptake | Float | 0, 25, 50, 75, 100 |
| max Phosphate uptake | Float | 0, 25, 50, 75, 100 |
| max Sulfur uptake | Float | 0, 25, 50, 75, 100 |
| max Oxygen uptake | Float | 0, 25, 50, 75, 100 |

**Table 11.**
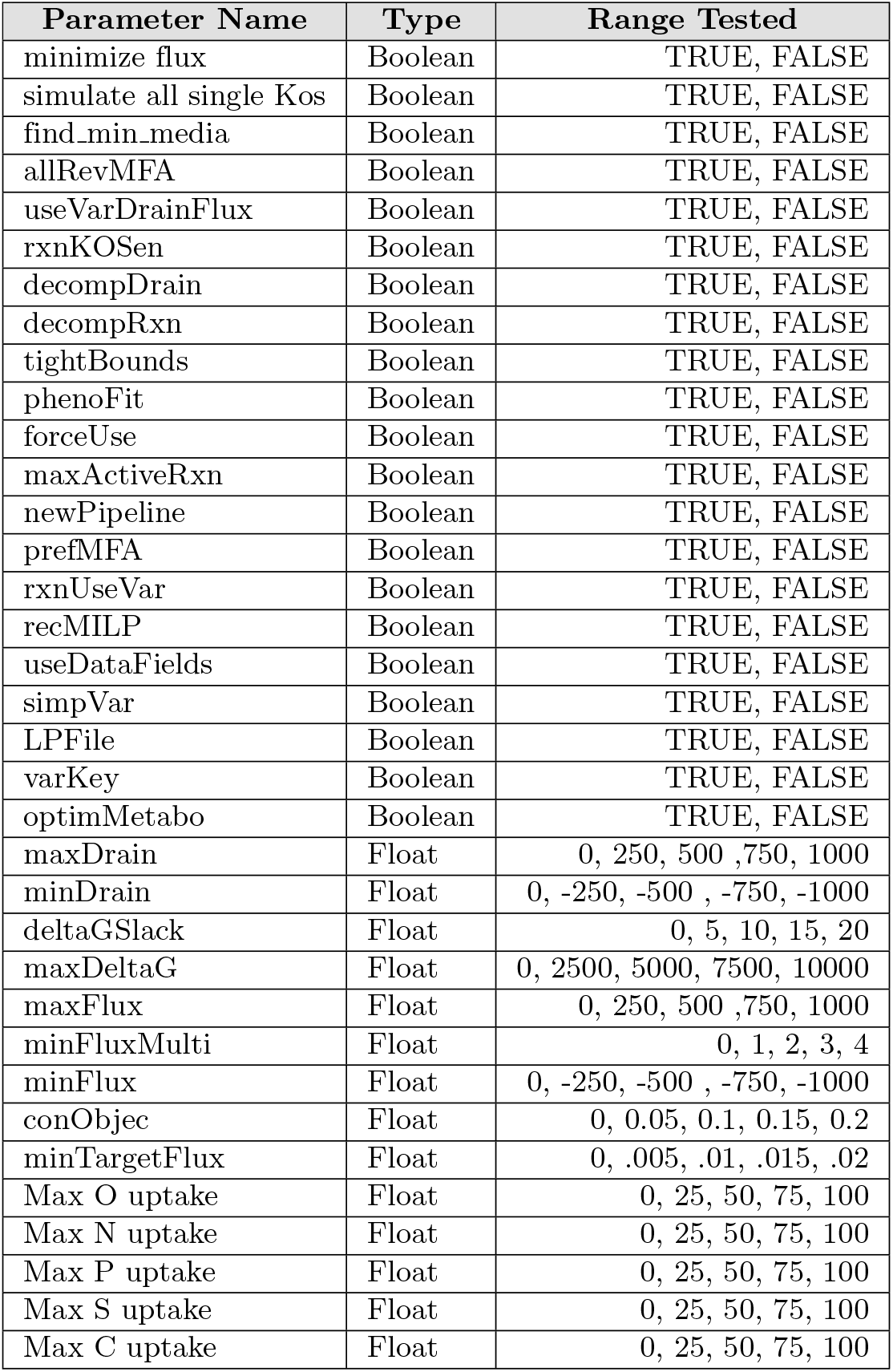
FBA-CLI configuration model

| Parameter Name | Type | Range Tested |
| --- | --- | --- |
| minimize flux | Boolean | TRUE, FALSE |
| simulate all single Kos | Boolean | TRUE, FALSE |
| find_min_media | Boolean | TRUE, FALSE |
| allRevMFA | Boolean | TRUE, FALSE |
| useVarDrainFlux | Boolean | TRUE, FALSE |
| rxnKOSen | Boolean | TRUE, FALSE |
| decompDrain | Boolean | TRUE, FALSE |
| decompRxn | Boolean | TRUE, FALSE |
| tightBounds | Boolean | TRUE, FALSE |
| phenoFit | Boolean | TRUE, FALSE |
| forceUse | Boolean | TRUE, FALSE |
| maxActiveRxn | Boolean | TRUE, FALSE |
| newPipeline | Boolean | TRUE, FALSE |
| prefMFA | Boolean | TRUE, FALSE |
| rxnUseVar | Boolean | TRUE, FALSE |
| recMILP | Boolean | TRUE, FALSE |
| useDataFields | Boolean | TRUE, FALSE |
| simpVar | Boolean | TRUE, FALSE |
| LPFile | Boolean | TRUE, FALSE |
| varKey | Boolean | TRUE, FALSE |
| optimMetabo | Boolean | TRUE, FALSE |
| maxDrain | Float | 0, 250, 500, 750, 1000 |
| minDrain | Float | 0, -250, -500, -750, -1000 |
| deltaGSlack | Float | 0, 5, 10, 15, 20 |
| maxDeltaG | Float | 0, 2500, 5000, 7500, 10000 |
| maxFlux | Float | 0, 250, 500, 750, 1000 |
| minFluxMulti | Float | 0, 1, 2, 3, 4 |
| minFlux | Float | 0, -250, -500, -750, -1000 |
| conObjec | Float | 0, 0.05, 0.1, 0.15, 0.2 |
| minTargetFlux | Float | 0, .005, .01, .015, .02 |
| Max O uptake | Float | 0, 25, 50, 75, 100 |
| Max N uptake | Float | 0, 25, 50, 75, 100 |
| Max P uptake | Float | 0, 25, 50, 75, 100 |
| Max S uptake | Float | 0, 25, 50, 75, 100 |
| Max C uptake | Float | 0, 25, 50, 75, 100 |

## Acknowledgments

This work has been supported in part by the National Science Foundation grant #CCF-1909688. The opinions and views presented are that of the authors and not of the NSF. This research has also been supported by the Plant-Microbe Interfaces Scientific Focus Area in the Genomic Science Program of the Office of Biological and Environmental Research (BER) at the U.S. Department of Energy Office of Science under Contract number DE-AC05-00OR22725. This research used resources of the Oak Ridge Leadership Computing Facility, which is a DOE Office of Science User Facility supported under Contract DE-AC05-00OR22725.

## Footnotes

1 We provide a publicly accessible KBase narrative demonstrating this example: https://doi.org/10.25982/135473.103/1960315

2 Released upon publication

3 Released upon publication

## References

1. Finkel OM, Salas-González I, Castrillo G, Conway JM, Law TF, Teixeira PJPL, et al. A single bacterial genus maintains root growth in a complex microbiome. Nature. 2020;587(7832):103–108.

2. Voit EO. Perspective: Dimensions of the scientific method. PLOS Computational Biology. 2019;15(9):1–14. doi:10.1371/journal.pcbi.1007279.

3. Sun SL, Yang WL, Fang WW, Zhao YX, Guo L, Dai YJ, et al. The Plant Growth-Promoting Rhizobacterium Variovorax boronicumulans CGMCC 4969 Regulates the Level of Indole-3-Acetic Acid Synthesized from Indole-3-Acetonitrile. Applied and Environmental Microbiology. 2018;84(16):e00298–18. doi:10.1128/AEM.00298-18.

4. Altschul SF, Gish W, Miller W, Myers EW, Lipman DJ. Basic local alignment search tool. Journal of molecular biology. 1990;215(3):403–410.

5. Shah N, Nute MG, Warnow T, Pop M. Misunderstood parameter of NCBI BLAST impacts the correctness of bioinformatics workflows. Bioinformatics. 2018;35(9):1613–1614. doi:10.1093/bioinformatics/bty833.

6. Bethesda (MD): National Center for Biotechnology Information (US). BLAST Command Line Applications User Manual; 2008. Available from: https://www.ncbi.nlm.nih.gov/books/NBK279690/.

7. Madden T, Busby B, Ye J. Reply to the paper: Misunderstood parameters of NCBI BLAST impacts the correctness of bioinformatics workflows. Bioinformatics. 2019;35(15):2699–2700.

8. González-Pech RA, Stephens TG, Chan CX. Commonly misunderstood parameters of NCBI BLAST and important considerations for users. Bioinformatics. 2018;35(15):2697–2698. doi:10.1093/bioinformatics/bty1018.

9. Morrison-Smith S, Boucher C, Bunt A, Ruiz J. Elucidating the role and use of bioinformatics software in life science research. In: Proceedings of the British HCI Conference. ACM; 2015. p. 230–238.

10. Jamshidi P, Siegmund N, Velez M, Kästner C, Patel A, Agarwal Y. Transfer Learning for Performance Modeling of Configurable Systems: An Exploratory Analysis. In: IEEE/ACM International Conference on Automated Software Engineering (ASE). IEEE; 2017. p. 497–508.

11. Cohen MB, Dwyer MB, Shi J. Constructing Interaction Test Suites for Highly-Configurable Systems in the Presence of Constraints: A Greedy Approach. IEEE Transactions on Software Engineering. 2008;34(5):633–650.

12. Miller G. Scientific publishing. A scientist’s nightmare: software problem leads to five retractions. Science (New York, NY). 2006;314(5807):1856.

13. Ichikawa T, Suzuki Y, Czaja I, Schommer C, Leßnick A, Schell J, et al. Retraction Note to: Identification and role of adenylyl cyclase in auxin signalling in higher plants. Nature. 1998;396. doi:10.1038/24659.

14. Orth JD, Thiele I, Palsson BØ. What is flux balance analysis? Nature Biotechnology. 2010;28(3):1546–1696.

15. FAQ: Metabolic Modeling; 2022. Available from: https://docs.kbase.us/workflows/metabolic-models/faq-metabolic-modeling.

16. Jin D, Qu X, Cohen MB, Robinson B. Configurations Everywhere: Implications for Testing and Debugging in Practice. In: International Conference on Software Engineering, Software in Practice Track. ICSE. ACM; 2014. p. 215–225.

17. Sayagh M, Hassan AE. ConfigMiner: Identifying the Appropriate Configuration Options for Config-related User Questions by Mining Online Forums. IEEE Transactions on Software Engineering. 2020; p. 1–1. doi:10.1109/TSE.2020.2973997.

18. Clements P, Northrop L. Software Product Lines: Practices and Patterns. Addison-Wesley Professional; 2001.

19. Johnson M, Zaretskaya I, Raytselis Y, Merezhuk Y, McGinnis S, Madden TL. NCBI BLAST: a better web interface. Nucleic Acids Research. 2008;36(2):W5–W9. doi:10.1093/nar/gkn201.

20. Yilmaz C, Cohen MB, Porter AA. Covering Arrays for Efficient Fault Characterization in Complex Configuration Spaces. IEEE Trans Software Eng. 2006;32(1):20–34. doi:10.1109/TSE.2006.8.

21. Zhang S, Ernst MD. Which Configuration Option Should I Change? In: Proceedings of the 36th International Conference on Software Engineering. ICSE 2014. New York, NY, USA: Association for Computing Machinery; 2014. p. 152–163. Available from: https://doi.org/10.1145/2568225.2568251.

22. Qu X, Cohen MB, Rothermel G. Configuration-aware Regression Testing: An Empirical Study of Sampling and Prioritization. In: International Symposium on Software Testing and Analysis. ISSTA. ACM; 2008. p. 75–86.

23. Wooley JC, Godzik A, Friedberg I. A primer on metagenomics. PLoS computational biology. 2010;6(2):e1000667.

24. Vollmers J, Wiegand S, Kaster AK. Comparing and evaluating metagenome assembly tools from a microbiologist’s perspective-not only size matters! PloS one. 2017;12(1):e0169662.

25. Smith TF, Waterman MS, et al. Identification of common molecular subsequences. Journal of molecular biology. 1981;147(1):195–197.

26. Kanehisa M, Goto S. KEGG: kyoto encyclopedia of genes and genomes. Nucleic acids research. 2000;28(1):27–30.

27. Cashman M, Cohen MB, Ranjan P, Cottingham RW. Navigating the Maze: The Impact of Configurability in Bioinformatics Software. In: Proceedings of the 33rd ACM/IEEE International Conference on Automated Software Engineering. ASE 2018. New York, NY, USA: ACM; 2018. p. 757–767.

28. Arkin AP, Cottingham RW, Henry CS, Harris NL, Stevens RL, Maslov S, et al. KBase: the United States department of energy systems biology knowledgebase. Nature biotechnology. 2018;36(7):566.

29. Altschul SF, Madden TL, Schäffer AA, Zhang J, Zhang Z, Miller W, et al. Gapped BLAST and PSI-BLAST: a new generation of protein database search programs. Nucleic acids research. 1997;25(17):3389–3402.

30. Camacho C, Coulouris G, Avagyan V, Ma N, Papadopoulos J, Bealer K, et al. BLAST+: architecture and applications. BMC bioinformatics. 2009;10(1):421.

31. Li D, Liu CM, Luo R, Sadakane K, Lam TW. MEGAHIT: an ultra-fast single-node solution for large and complex metagenomics assembly via succinct de Bruijn graph. Bioinformatics. 2015;31(10).

32. Li D, Luo R, Liu CM, Leung CM, Ting HF, Sadakane K, et al. MEGAHIT v1. 0: a fast and scalable metagenome assembler driven by advanced methodologies and community practices. Methods. 2016;102:3–11.

33. KBase MEGAHIT SDK Repository; 2017. Available from: https://github.com/kbaseapps/kb_megahit.

34. Henry CS, DeJongh M, Best AA, Frybarger PM, Linsay B, Sevens RL. High-throughput generation, optimization and analysis of genome-scale metabolic models. Nature Biotechnology. 2010;28(9):977–982.

35. Henry CS. MFAToolkit GitHub Repository; 2017. Available from: https://github.com/cshenry/fba_tools/tree/master/MFAToolkit.

36. Wilkinson MD, Dumontier M, Aalbersberg IJ, Appleton G, Axton M, Baak A, et al. The FAIR Guiding Principles for scientific data management and stewardship. Scientific data. 2016;3(1):1–9.

37. Lamprecht AL, Garcia L, Kuzak M, Martinez C, Arcila R, Martin Del Pico E, et al. Towards FAIR principles for research software. Data Science. 2020;3(1):37–59.

38. Shade A, Teal TK. Computing workflows for biologists: a roadmap. PLoS biology. 2015;13(11):e1002303.

39. Bechhofer S, De Roure D, Gamble M, Goble C, Buchan I. Research objects: Towards exchange and reuse of digital knowledge. Nature Precedings. 2010; p. 1–1.

